# Ecological niche divergence or ecological niche partitioning in a widespread Neotropical bird lineage

**DOI:** 10.1101/2024.01.18.576294

**Authors:** Jacob C. Cooper

## Abstract

Ecological niche divergence is generally considered to be a facet of evolution that may accompany geographic isolation and diversification in allopatry, contributing to species’ evolutionary distinctiveness through time. The null expectation for any two diverging species in geographic isolation is that of niche conservatism, wherein populations do not rapidly shift to or adapt to novel environments. Here, I test ecological niche divergence for a widespread, pan-American lineage, the avian genus of martins (*Progne*). Despite containing species with distributions that go from continent-spanning to locally endemic, I found limited evidence for niche divergence across the breeding distributions of *Progne*, and much stronger support for niche conservatism with patterns of niche partitioning. The ancestral *Progne* had a relatively broad ecological niche, similar to extant basal *Progne* lineages, and several geographically localized descendant species occupy only portions of the larger ancestral *Progne* niche. I recovered strong evidence of breeding niche divergence for four of 36 taxon pairs but only one of these divergent pairs involved two widespread, continental species (Southern Martin *P. elegans* vs. Gray-breasted Martin *P. chalybea*). Potential niche expansion from the ancestral species was observed in the most wide-ranging present-day species, namely the North American Purple Martin *P. subis* and *P. chalybea*. I analyzed populations of *P. subis* separately, as a microcosm of *Progne* evolution, and again found only limited evidence of niche divergence. This study adds to the mounting evidence for niche conservatism as a dominant feature of diversifying lineages. Even taxa that appear unique in terms of habitat or behavior may still not be diversifying with respect to their ecological niches, but merely partitioning ancestral niches among descendant taxa.

## 1 Introduction

Species’ ecological niches, like morphological traits, evolve via natural selection, thus altering their ecological niches and geographic potential through time (Engler et al., 2021). These processes are particularly important for diversification among closely related species, as niche divergence and diversification can lead to or reinforce speciation (Hu et al., 2015; Cuervo et al., 2021; Şahin et al., 2021). Such ecological shifts can lead to dramatic shifts in geographic distributional potential, and may follow predictable patterns through evolutionary time (Cobos et al., 2021). For example, insular species often shift from lowland to montane situations through time (Ricklefs & Cox, 1972; Kennedy et al., 2022), whereas the opposite tendency (highlands to lowlands) may dominate in continental settings (van Els et al., 2021).

Despite frequent opportunities for species to adapt and evolve with respect to their ecological niches, ecological niche conservatism appears to be the norm within most species (Peterson, Soberón & Sanchez-Cordero, 1999; Peterson, 2011; Khaliq et al., 2015; García-Navas & Westerman, 2018), at least with respect to coarse-resolution environmental conditions (Comte, Cucherousset & Olden, 2017). Such conservatism has been argued to be a contributing factor in diversification dynamics, especially in systems in which conserved niches through time force populations into allopatry and allow independent evolution to occur (Prigogine, 1987; Vrba, 1993; Kozak & Wiens, 2006). Indeed, in birds, secondary contact is often identified as a driving force for character divergence, including in ecological niches (Endler, 1977; Seddon & Tobias, 2007; McCormack, Zellmer & Knowles, 2009).

Three major scenarios for niche evolution are thus available for allopatric and parapatric populations sharing a recent common ancestor: (1) descendant populations become wholly allopatric and undergo no appreciable niche differentiation; (2) descendant populations occupy different parts of their ancestor’s ecological niche in allopatry or parapatry, adapting to these specific conditions and partitioning ecological space upon subsequent secondary contact; and (3) one or more descendant populations are able to adapt to new environments and occupy novel ecological niches (Figure 1). Scenario 1 appears to be the norm, but these patterns may be overridden in deeper time by the open exchange of genes between populations after secondary contact, as has occurred with raven lineages (*Corvus* spp.) in North America (Omland, Baker & Peters, 2006). Scenario 2 is perhaps best exemplified by *Poecile* chickadees within North America: two lineages (Carolina Chickadee *P. carolinensis* and Black-capped Chickadee *P. atricapillus*) have a narrow but distinct hybrid zone within which each species can survive, with the socially dominant *P. carolinensis* slowly pushing northwards as the climate warms (Mostrom, Curry & Lohr, 2020). These situations can also lead to complicated hybrid zones, as in taxa that now exist in secondary contact after retreating to different Pleistocene refugia (*e.g.*, members of the Yellow-rumped Warbler *Setophaga coronata* complex) (Hubbard, 1969; Milá, Smith & Wayne, 2007). Scenario 3 is perhaps less frequent, but is likely manifested within genera such as *Baeolophus* titmice, in which the interior western Juniper Titmouse *Baeolophus ridgwayi* exists in drier, more xeric conditions than all of its congeners (Cicero, Pyle & Patten, 2020).

**Figure 1.**
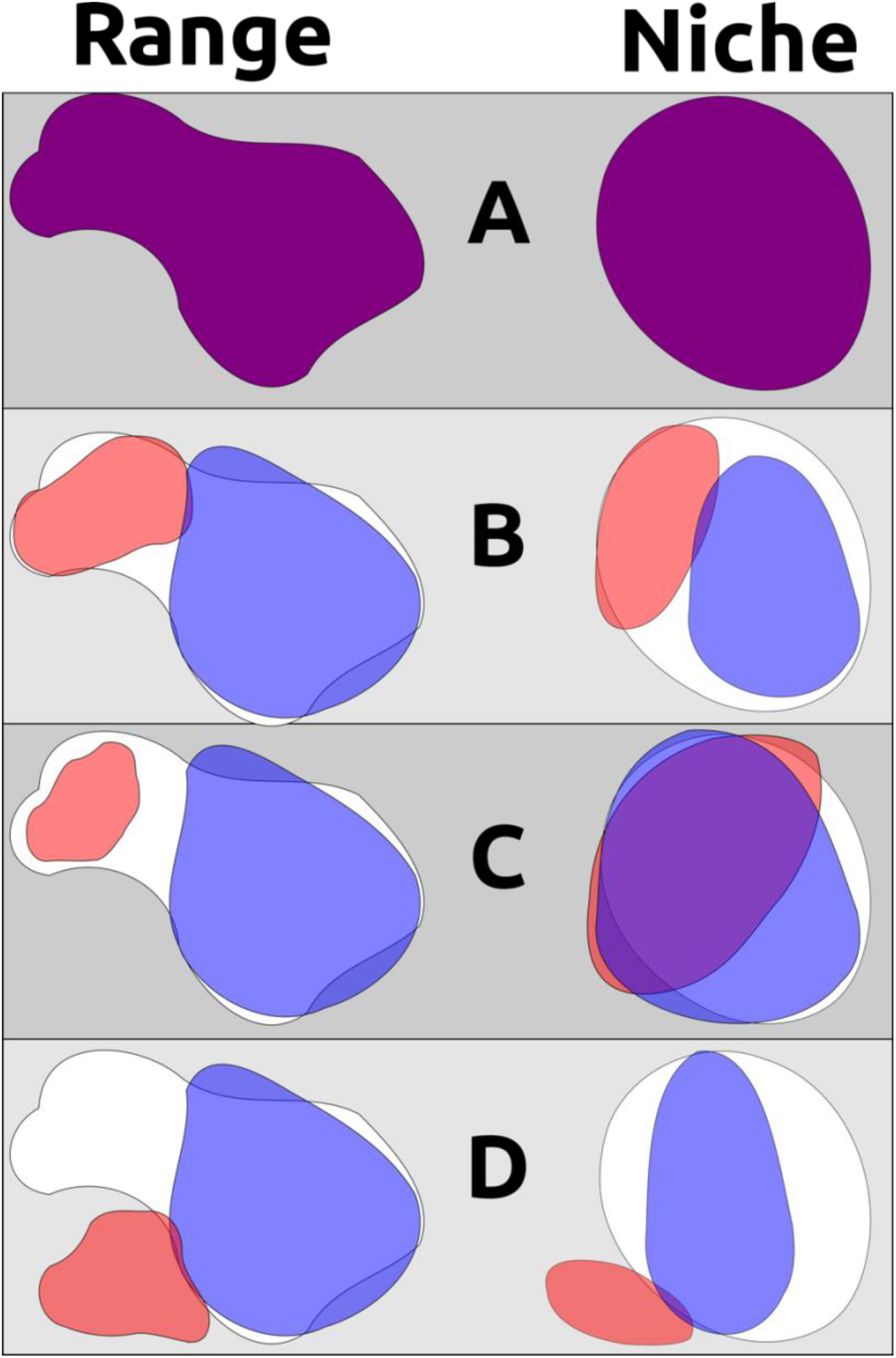
A non-exhaustive demonstration of different ways geographic range and ecological niche can evolve independently. A) an example ancestral species’ range and ecological niche, with outlines of these characters shown in the background for subsequent descendant examples; B) two descendant populations partition the ancestor’s geographic range and ecological niche, such as when a pan-archipelagic species diversifies within individual island groups into discrete ranges and niches that are different from each other but not from the ancestral species (*e.g.*, *P. cryptoleuca* and *P. dominicensis*); C) two descendant populations occurring in allopatry but occupying non-differentiable ecological niches and exhibiting niche conservatism; D) two descendant populations where one occupies a portion of the ancestral range and ancestral niche while the other other has evolved to occupy a new range and new niche (assuming that the ancestral species was occupying the entirety of its potential geographic distribution and the entirety of its ecological niche). Note that B and C are indicative of niche conservatism through time, even though pairwise tests of B may show ecological niche divergence in the present day.

One genus that likely has undergone multiple modes of diversification is the martin genus *Progne* (Aves: Hirundinidae) (Table 1). These aerial insectivores are distributed from central Canada to southern Argentina, nesting in cavities in trees, rocks, man-made structures, and even occasionally in the ground (Allen & Nice, 1952; Pistorius, 1975). Northern and southern representatives of the genus are migratory, with several resident taxa overlapping with these species in the Tropics during non-breeding periods. Movements of *Progne* species appear to be complex, with some species undergoing seasonal inter-regional movements between primary and secondary wintering areas thousands of kilometers apart (Siddiqui, 2017). Even daily movements of *Progne* can cover large distances, with coastal, sea-level-nesting Peruvian Martins *P. murphyi* reaching elevations as high as 1500 m (Parker, Stotz & Fitzpatrick, 1996; Luo, 2020) and Purple Martins *P. subis* foraging as high as 1889 m above the ground (Helms et al., 2016), sometimes quite far from nest sites (Corman, 2005). The combination of aerial feeding habits and migratory behaviors enables *Progne* to cover long distances as migrants and as vagrants, as demonstrated by records *P. subis*, a species that breeds broadly across central North America and winters in central South America, from Alaska, Ireland, Scotland, the Azores, and the Falklands [Malvinas] (eBird, 2012; Quigley, 2018; Brown, Airola & Tarof, 2021).

**Table 1.**
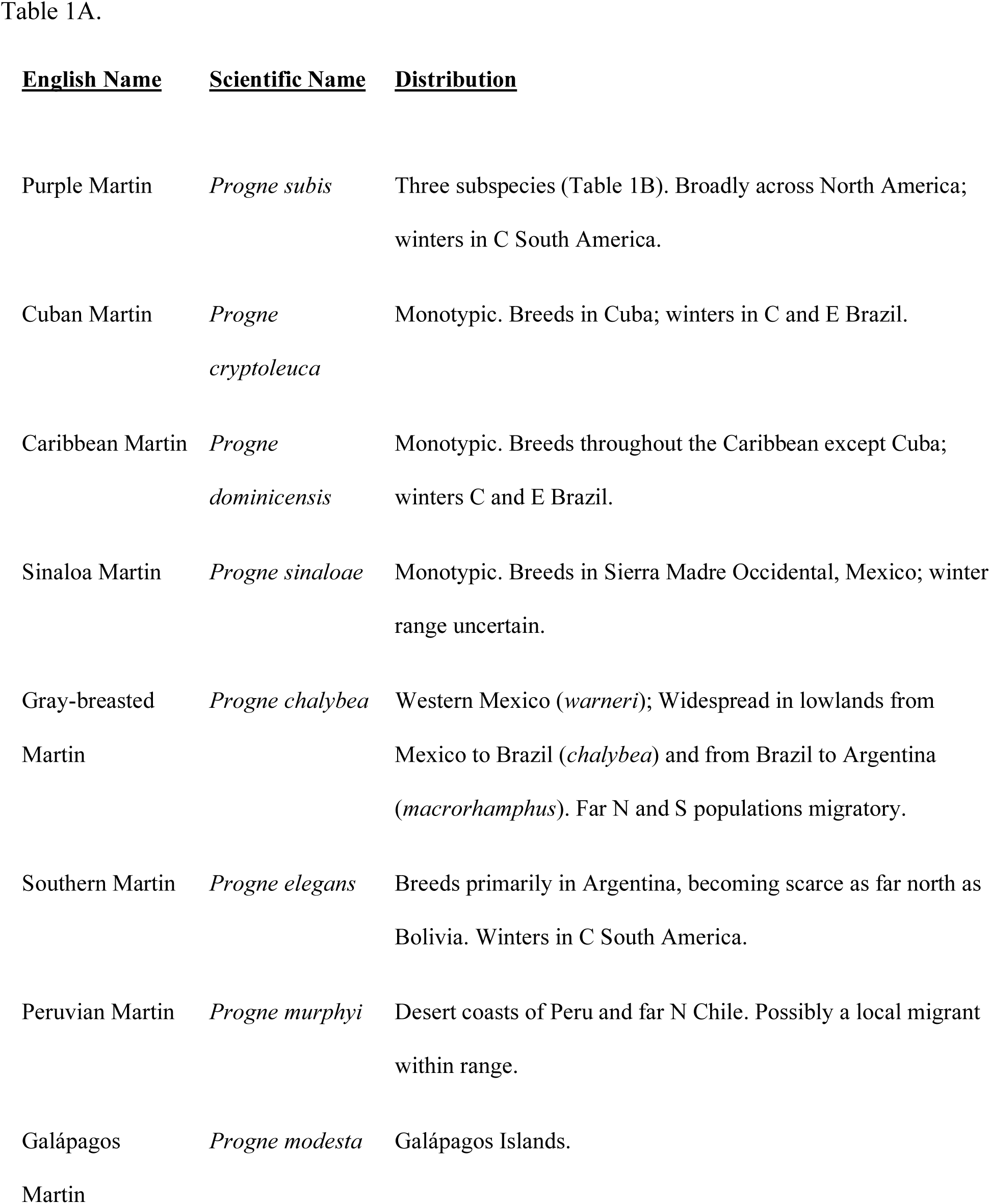
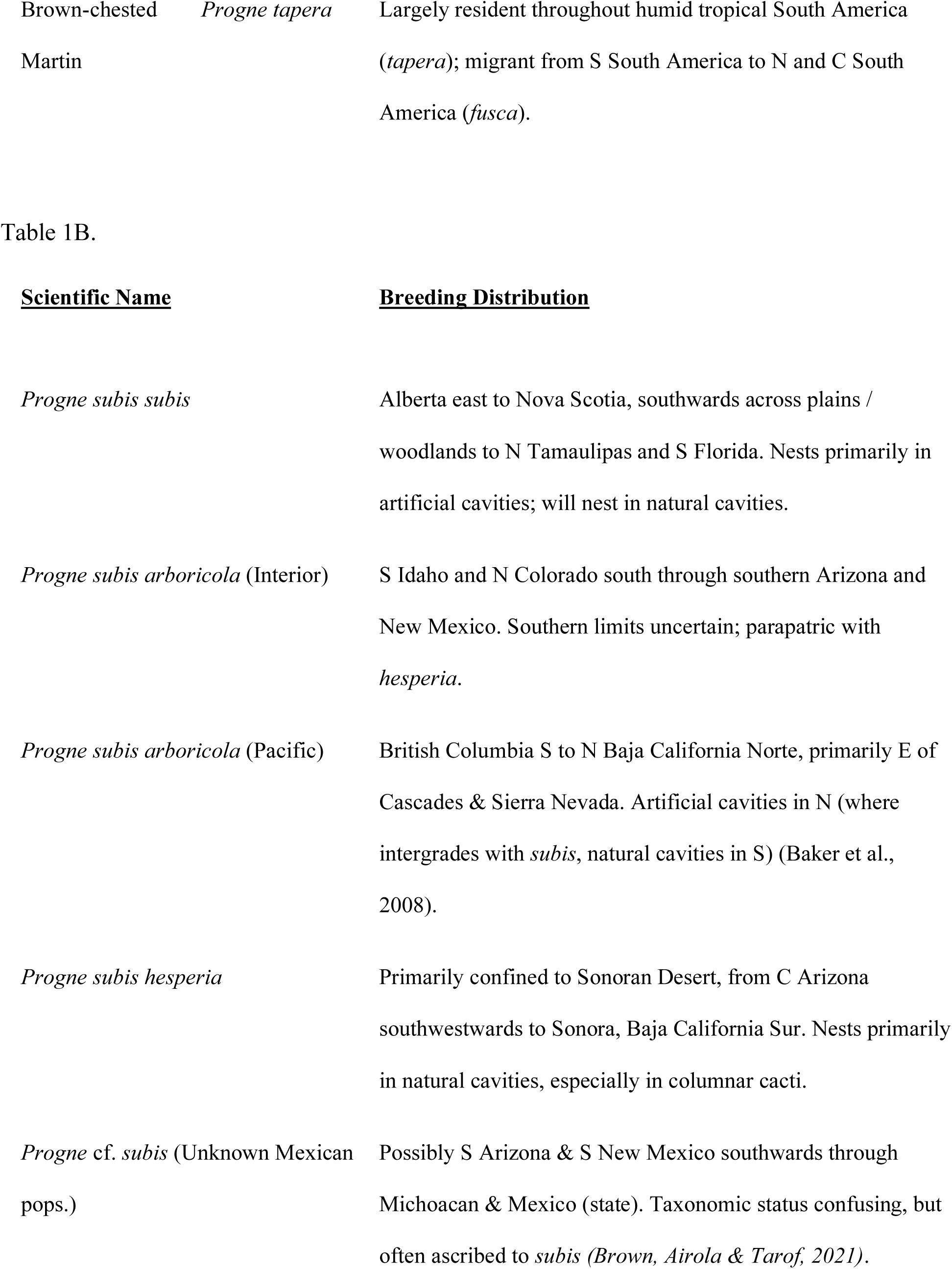
*Part A*: a list of all *Progne* species with subspecific breakdown for all taxa except *subis*. *Part B*: distributions of study populations within *P. subis* used in this paper and their respective subspecific groups.

Martins show high levels of phenotypic conservatism, such that winter distributions for many species are unknown or are only being confirmed in recent years; many migrant or away-from-breeding-range individuals (especially among the ‘white-breasted’ martin complex of the Caribbean and Mexico) are not identifiable in the field (Fang & Schulenberg, 2020; García-Lau & Turner, 2021; Perlut & Williams, 2021). Distributions vary greatly in extent: the Galápagos Martin *P. modesta (Roper, 2020)* and *P. murphyi (Luo, 2020)* inhabit geographically limited xeric coastal areas whereas most widespread species are found across continents either seasonally or year round. An extreme example is the Gray-breasted Martin *P. chalybea*, which breeds across the entire latitudinal breadth of the Tropics from Mexico to Argentina (Lagasse, 2020).

Given that this genus is monophyletic and its phylogenetic relationships have been well documented (Moyle et al., 2008; Brown, 2019), it is possible to explore the evolutionary history of ecological niche traits. Studying ecological niche evolution in *Progne* provides an opportunity to examine niche differentiation in a wide-ranging genus that has evolved migratory and locally endemic resident forms that are largely conserved morphologically, as well as to test whether a geographically widespread species complex has evolved to take advantage of new ecological niches or has merely partitioned the ancestral species’ ecological niche. Martin species occur in allopatry, parapatry, and sympatry in their breeding distributions, allowing for tests of niche conservatism between pairs of species that are of varying relatedness and that possess varying levels of geographic overlap. Reconstructing niche evolution in this genus will also shed light on what kinds of evolutionary shifts lead to local endemism, and will provide a continental comparison to island speciation cycles.

## 2 Methods

### 2.1 General Data Cleaning

Occurrence data for all *Progne* martin species were downloaded from the Global Biodiversity Information Facility (GBIF) on 21 Feb 2022 (Global Biodiversity Information Facility, 2022). Data were processed using R 4.1.2, 4.2.0, and 4.2.1 (R Core Team, 2022) , relying on the general data packages of *tidyverse* (Wickham et al., 2019) and *data.table (Dowle & Srinivasan, 2019)*, and the general spatial packages *maptools* (Bivand & Lewin-Koh, 2022)*, raster* (Hijmans, 2022), *rgdal* (Bivand, Keitt & Rowlingson, 2022), and *sf* (Pebesma, 2018). Data were reduced to presence-absence data based at localities, with duplicate sightings of particular species from single sites removed. Distributions of individual species were superimposed on country borders using *rnaturalearth* (South, 2017) and *rnaturalearthhires* (South, 2022), and occurrences of each species were compared to known distributions for each taxon (Billerman et al., 2020). Outlier occurrences for each taxon were removed or re-identified depending on their location, the taxonomic history for the species in question, and on the species’ residency status in a given region (see Supplemental Materials). These steps are necessary given the number of misidentifications and low-confidence identifications in the database at the time of download, and given the long-distance dispersal ability of *Progne*. Based on published distributions and breeding records, I created dispersal areas by hand based on major biogeographic barriers (e.g., straights, crests of mountains) referred to as **M**s for identifying potentially erroneous points and, for allopatric complexes like *P. subis*, assigning breeding records to the correct geographic populations (Soberón & Peterson, 2005; Cooper et al., 2021). I also ran a custom ‘rarefy’ code provided by Dr. J. D. Manthey (Texas Tech University) to thin occurrences to be ≥20 km to one another using the *R* package *fossil* (Vavrek, 2011). This last step was particularly important near large population centers or known colonies for rarer taxa, where records can be much denser than rural areas, potentially introducing biases in the models.

Occurrence data were largely restricted to April-July (Northern Hemisphere) and October-January (Southern Hemisphere) for migratory taxa to compare breeding niches, with minor adjustments for individual species based on their migratory patterns (see Supplemental Material). I focused on summer distributions as winter distributions are less well-known and are data-depauperate, especially for long-distance migrants. These restrictions ensured that comparisons of taxa encompassed similar phenological periods, and that records could be identified with greater confidence, as species’ ranges are known to be largely discrete in breeding season. Non-breeding distributions of many migratory taxa remain incompletely known, such that whether non-breeding distributions of several species pairs are wholly syntopic is unknown (Perlut, Klak & Rakhimberdiev, 2017; Fang & Schulenberg, 2020; Turner, 2020; Brown, Airola & Tarof, 2021; García-Lau et al., 2021; García-Lau & Turner, 2021; Perlut & Williams, 2021).

These steps were also applied to subspecies of *Progne subis* for analyses within a polytypic migratory taxon. I did not repeat these analyses with Brown-chested Martin *P. tapera*, the other polytypic *Progne* with migratory populations, as the distributions of migratory and non-migratory populations do not appear to be discrete, and many records within the GBIF database are not identified to subspecies. For *Progne subis*, three subspecies are described, but the limits of their distributions are not well defined (Brown, Airola & Tarof, 2021). Specifically, among western populations, lines of evidence for subspecies assignment, behaviors, and nesting preferences vary greatly across the species’ range. As such, I subdivided *P. subis* into the following populations for analysis: nominate *P. s. subis* of eastern North America; core *P. s. arboricola* in the Rocky Mountains as far south as the Mogollon Rim; *P. s. hesperia* of the Sonoran desert; Pacific coast *P. s. arboricola* of California north to British Columbia; and interior Mexican populations of unknown taxonomic status from the mountains south of the Mogollon Rim to southwestern Mexico (Brown, Airola & Tarof, 2021).

### 2.2 Environmental Data

Environmental data were drawn from the ENVIREM dataset (Title & Bemmels, 2018). I removed count-format data restricting the data to continuous, raster-format variables representing terrestrial conditions. I retained elevation in the analyses but I removed the terrain roughness index, as elevation is known to affect the physiology of birds (and thus may affect nest site selection) but general terrain roughness is likely to affect *Progne* only indirectly, given that single species can be found under very diverse topographic conditions (Dubay & Witt, 2014). The remaining environmental variables were transformed via principal components analyses using the function ‘rda’ in the R package *vegan* (Oksanen et al., 2022). The first principal component was mostly explained by variation in potential evapotranspiration (96.3%), annual potential evapotranspiration (1.2%) and thermicity (a metric comparing minimum and average temperatures; 1.2%). The second principal component was likewise dependent on evapotranspiration, with the primary explaining variables of annual evapotranspiration (70.1%), thermicity (18.7%), and maximum temperature of the coldest month (5.7%).

Data were further restricted to variables less affected by temperate latitude seasonality to reflect potential differences within populations’ breeding niches better. That is, I retained the ENVIREM variables of annual potential evapotranspiration, Thornthwaite aridity index, climatic moisture index, Emberger’s pluviometric quotient (a measure for differentiating Mediterranean climates), minimum temperature of the warmest month (generally corresponding with the breeding season for migratory taxa), potential evapotranspiration of the driest quarter, potential evapotranspiration of the warmest quarter, potential evapotranspiration of the wettest quarter, and topographic wetness index (Title & Bemmels, 2018). The top remaining explanatory variables for the first principal component were annual evapotranspiration (70.3%) and Emberger’s pluviometric quotient (27.8%). For the second principal component, they were Emberger’s pluviometric quotient (71.5%) and annual evapotranspiration (27.3%). I only used the first two principal components to analyze variable importance and to look at niche differentiation, and I used variables directly for creating ecological niche models. I proceeded with all subsequent analyses using this subset of environmental data.

### 2.3 Environmental Comparisons

To assess different ‘ecopopulations’ (*i.e.*, populations as defined by unique environments) and to identify how well-partitioned *Progne* taxa are ecologically, I performed linear discriminant analyses in *R*, using the ‘lda’ function in the package *MASS* (Venables & Ripley, 2002). These tests partition individuals based on the environmental data, and suggest hypotheses for group assignments for individuals among known group assignments and groups presumed based on environmental characteristics (Cooper et al., 2021). I performed these tests both for all *Progne* species and within *P. subis*. I also verified whether the number of taxa recognized is supported by the environmental data by performing gap-statistic analyses of *k*-means clusters using the function ‘fviz_nbclust’ in the *R* package *factoextra (Kassambara & Mundt, 2020)*.

### 2.4 Niche Modeling and Comparisons

For each species, I created ecological niche models using a presence-only method, minimum volume ellipsoids, following Cooper et al. (2021). Specifically, I fit minimum volume ellipsoids with an inclusion level of 90% using the ‘cov.mve’ function in the *R* package *MASS* (Venables & Ripley, 2002). I departed from the previous pipeline with respect to thresholding to create a presence-absence map. Whereas this threshold was previously determined by applying a normal distribution cutoff to the Mahalanobis distances within the data (Cooper et al., 2021), these data are often non-normal, and such thresholding can augment bias in the dataset. These distances are better fit by a Gaussian distribution, given the overwhelming number of points close to the centroid and the right skew of the data. Gaussian distributions fitting each ellipsoid were determined using the *R* function ‘fitdist’ in the package *fitdistrplus* (Delignette-Muller & Dutang, 2015); models were then thresholded to the following inclusion levels: 75%, 85%, 90%, 95%, and 99%. These models were run for every *Progne* species, as well as for subpopulations of *P. subis*.

I performed pairwise niche comparisons of species following Warren et al. (2008) and expanded by Cooper et al. (2021) utilizing the comparative statistic of Schoener’s *D* calculated using the R package *dismo* (Hijmans et al., 2021). Schoener’s *D* returns a value between 0 and 1, with 0 being wholly different ecological niche models and 1 being identical ecological niche models. For each pair of species *A* and *B*, I computed the observed value of Schoener’s *D* for *A* and *B*, and for two test distributions derived from 100 ecological niche models created from random points within each population’s accessible area (**M**) (Warren, Glor & Turelli, 2008; Glor & Warren, 2011; Cooper et al., 2021). Specifically, I defined a test statistic as the ‘true’ comparison between *A* and *B* using the species’ ecological niche models, and then derived the two random distributions by comparing the model of *A* to the models derived from random points from across the accessible area (**M**) of *B* and *vice versa* (Glor & Warren, 2011; Cooper et al., 2021). I considered tests above the 97.5% confidence interval for both random distributions to be a very clear failure to reject the null hypothesis, and clear confirmation that *A* and *B* should be considered to have highly similar, near-equivalent ecological niches (Glor & Warren, 2011; Cooper et al., 2021). If the statistic fell below the 2.5% confidence interval, *A* and *B* were assessed to have divergent ecological niches, and thus be a rejection of the null hypothesis. Many other outcomes are possible, most indicating that the contrast between is indistinguishable from random, or that the two tests are not both significant in the same direction. These tests could be indicative of a ‘spectrum’ of ongoing niche diversification, but I treat them separately; in the end, however, they are not rejections of the null hypothesis of ecological niche conservatism (Cooper et al., 2021).

### 2.5 Historical Niche Reconstruction

Historical niche reconstructions were performed using the R packages *ellipsenm* (Cobos et al., 2022), *geiger* (Pennell et al., 2014), *nodiv* (Borregaard et al., 2014), and *nichevol* (Cobos, Owens & Peterson, 2020), taking into account accessible areas (**M**s) for each species and the lack of access of many species to all climatic combinations experienced by the genus (Saupe et al., 2018). I used the same ecological characters as described above for the pairwise comparisons for consistency (Cooper & Soberón, 2018). I used a phylogenetic tree of species’ relationships based on a UCE study of the family Hirundinidae provided by Clare E. Brown, Subir Shakya, and Fred Sheldon (Brown, 2019). This tree is missing two taxa due to poor DNA reads, namely Galápagos *Progne modesta* and Cuban *P. cryptoleuca* Martins. I added *P. cryptoleuca* to the tree as sister to Caribbean Martin *P. dominicensis* halfway between the node and the base of the dendrogram based on phylogenetic information indicating that these taxa form a sister-pair (Moyle et al., 2008). I performed *nichevol* analyses of each ENVIREM character to assess how ecological niches shifted through time, and to identify which species experience the largest evolutionary shifts. shifted through time, and to identify which species experience the largest evolutionary shifts.

### 2.6 General Note on Results

Many of the results presented herein involve models that are not static and depend on dynamic parameters; as a result, exact values may vary between iterations, but general patterns were consistent between all iterations I performed within this study.

## 3 Results

### 3.1 Environmental Differences and Clustering

I observed broad ecological niche overlap within the genus generally, with respect to individual environmental variables and in terms of environmental variables transformed by principal components analysis. Species that occupy different extreme habitats (*e.g.,* desert vs. rainforest) may show differentiation along individual environmental axes that reflect these differences, but few taxa showed overall differences in realized niches (see Supplementary Material). Using gap-statistic analysis, I determined the number of ‘ecospecies’ within the genus *Progne* to be 5 (see Supplemental Material). However, when classifying the full set of occurrence data into five groups using *k*-means, none of these ecospecies corresponded clearly to any described taxon, and no individual taxon is fully within any *k*-means group (Supplemental Material). Using discriminant function analyses, the most accurately reconstructed species from environmental data were *P. murphyi* (90% clustered together) and *P. subis* (94%). The greatest confusion was related to Gray-breasted Martin *P. chalybea*, a species that is wide-ranging both geographically and environmentally. Most individuals of Cuban Martin *P. cryptoleuca*, Caribbean Martin *P. dominicensis*, Sinaloa Martin *P. sinaloae*, and Brown-chested Martin *P. tapera*, were ascribed to the same group as *P. chalybea* (Table 2).

**Table 2.** Percent of individuals correctly clustered to original species using environmental data in a discriminant function analysis, with original (correct) taxon assignments being shown as rows and predicted taxon cluster being shown as columns, with accurate groupings falling along the bolded diagonal. Values shown are rounded percents and thus may not exactly add to 100, and taxa are listed alphabetically.

| ***Progne species*** | *chalybea* | *cryptoleuca* | *dominicensis* | *elegans* | *modesta* | *murphyi* | *sinaloae* | *subis* | *tapera* |
| --- | --- | --- | --- | --- | --- | --- | --- | --- | --- |
| *chalybea* | **78** | 0 | 2 | 0 | 0 | 2 | 0 | 6 | 11 |
| *cryptoleuca* | 94 | **0** | 6 | 0 | 0 | 0 | 0 | 0 | 0 |
| *dominicensis* | 62 | 0 | **29** | 0 | 9 | 0 | 0 | 0 | 0 |
| *elegans* | 9 | 0 | 0 | **53** | 0 | 0 | 0 | 27 | 11 |
| *modesta* | 0 | 0 | 25 | 0 | **75** | 0 | 0 | 0 | 0 |
| *murphyi* | 10 | 0 | 0 | 0 | 0 | **90** | 0 | 0 | 0 |
| *sinaloae* | 80 | 0 | 0 | 0 | 0 | 0 | **10** | 10 | 0 |
| *subis* | 1 | 0 | 0 | 3 | 0 | 0 | 0 | **94** | 1 |
| *tapera* | 51 | 0 | 1 | 3 | 0 | 0 | 0 | 21 | **22** |

Within *P. subis*, I found that the most-supported number of ecopopulations in this clade is 1. This result holds true even when all taxa except nominate *P. s. subis* are compared. Despite this, discriminant function analyses of the three currently recognized subspecies of *P. subis* had a high level of success in discriminating taxa, with accuracies of 96% for *P. s. arboricola*, 96% for *P. s. hesperia*, and 99% for *P. s. subis* (see Supplemental Materials). Subdividing these taxa further into the five groups based on region, habitat, and behavior still showed high support for each described subspecies, with accuracies of 93% for *P. s. arboricola*, 90% for *P. s. hesperia*, and 100% for *P. s. subis* (Table 3). Additionally, Pacific *P. s. arboricola* populations were largely differentiable from interior *arboricola* populations, with 82% of individuals being correctly identified and only 12% of individuals erroneously assigned as interior *arboricola*. Mexican populations were split between being considered their own entity (40%) and being considered an extension of interior *arboricola* (32%).

**Table 3.** Percent of individuals correctly clustered to original subspecies within *P. subis* using environmental data in a discriminant function analysis, with original (correct) taxon assignments being shown as rows and predicted taxon cluster being shown as columns, with accurate groupings falling along the bolded diagonal. Values shown are rounded percents and thus may not exactly add to 100, and taxa are listed alphabetically.

| *Progne subis*<br>Populations | *arboricola*<br>(Interior) | *arboricola*<br>(Pacific) | *hesperia* | *subis* | Mexican pops. |
| --- | --- | --- | --- | --- | --- |
| *arboricola*<br>(Interior) | **93** | 1 | 1 | 0 | 0 |
| *arboricola*<br>(Pacific) | 12 | **82** | 4 | 0 | 2 |
| *hesperia* | 4 | 1 | **90** | 0 | 4 |
| *subis* | 0 | 0 | 0 | **100** | 0 |
| Mexican pops. | 32 | 8 | 12 | 8 | **40** |

### 3.2 Ecological Niche Divergence

Niches appeared to be largely conserved within the genus *Progne*, with a failure to reject the null hypothesis in 69% of pairwise tests; significant failures to reject with respect to both comparisons were only found in two pairwise comparisons: *P. subis* vs. Southern Martin *P. elegans* and *P. subis* vs. *P. tapera*) (Table 4A; Figure 2). Only four pairwise tests showed definitive ecological niche divergence between test taxa: *P. chalybea* vs. *P. elegans, P. chalybea vs, P. murphyi*, *P. cryptoleuca* vs. *P. elegans*, and *P. cryptoleuca* vs. *P. murphyi*. Rejection of the null hypothesis for only one of the two pairwise comparisons was observed in six more comparisons, most involving combinations with *P. dominicensis*, *P. tapera*, *P. murphyi*, and *P. modesta* (Table 4A). Instances of divergence were not limited to widespread species or limited-range species, with one instance of divergence found between two wide-ranging species (*Progne chalybea* and *P. elegans*), two instances between widespread and restricted-range species (*P. chalybea* and *P. murphyi*, *P. elegans* and *P. cryptoleuca*), and one between two restricted-range species (*P. murphyi* and *P. cryptoleuca*). Conversely, ecological niche comparisons that were most similar (*i.e.,* the null of conservatism was the least likely to be rejected) involved the ecologically diverse *P. subis*. Among subpopulations of *Progne subis*, no comparisons were able to conclusively reject the null hypothesis of niche conservatism (Table 4B). Only two pairwise comparisons rejected the null hypothesis for one of the two comparisons: *P. s. arboricola* (Rocky Mountains) vs. *P. s.* unknown (Mexico), and *P. s. hesperia* vs. *P. s.* unknown (Mexico). The most similar taxa appeared to be *P. s. arboricola* (Pacific Coast) and both *P. s. arboricola* (Rocky Mountain) and *P. s. hesperia*; however, it is unclear whether this similarity is informative or merely within the variation of the spectrum of what can be considered niche conservatism.

**Figure 2.**
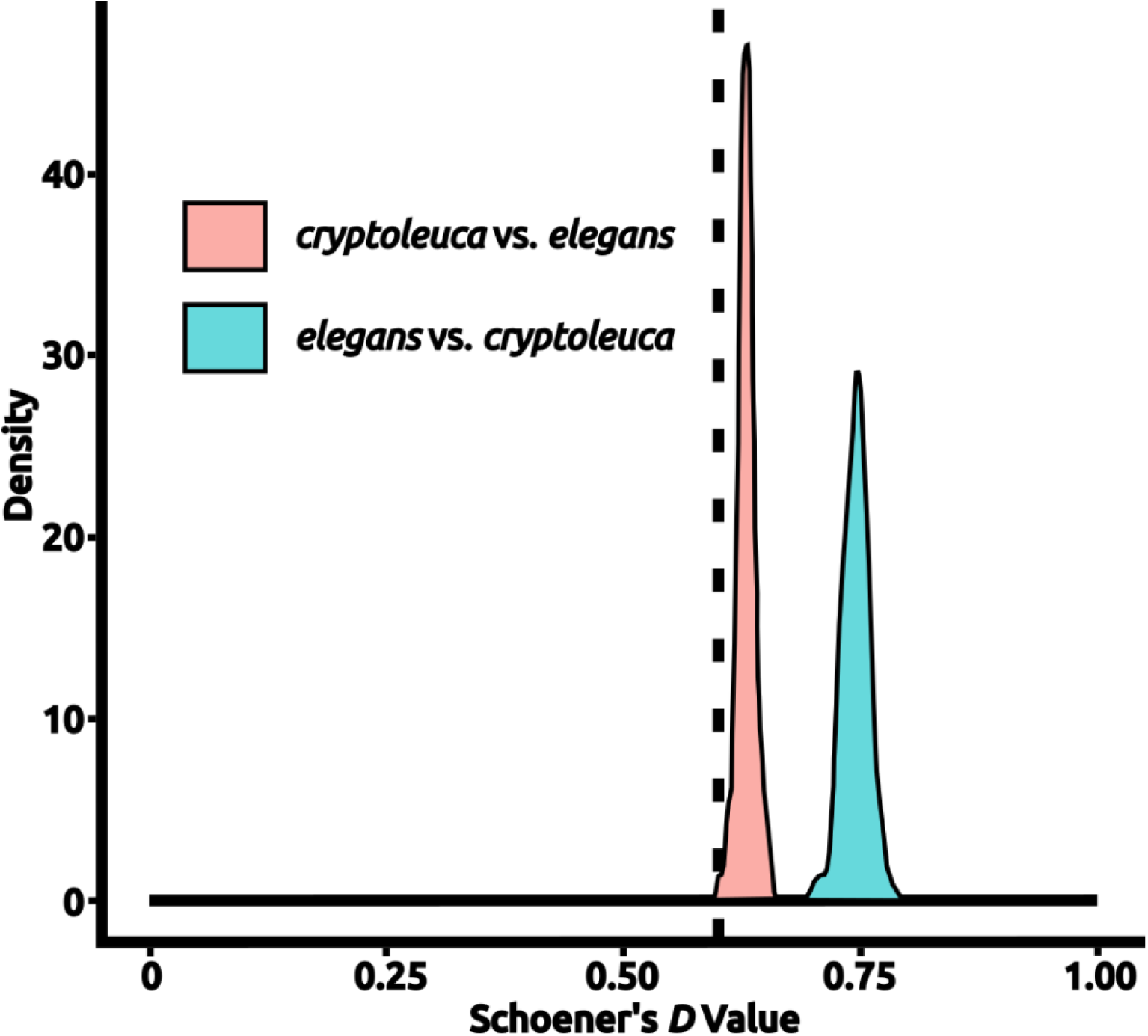
An example test of niche equivalency. Density histograms show the distribution of values drawn from calculating Schoener’s *D* for niche models created from random points in each species’ accessible area to the actual niche model created for the other species, with the test statistic being derived from a direct comparison of the “true” models created from each species’ occurrences. Values of Schoener’s *D* vary between 0 (completely different) and 1 (wholly identical). In this example, *Progne elegans* is compared to *P. cryptoleuca*, with a density histogram of values from random tests being shown; the test statistic is shown as a vertical dashed line. In this particular example, the taxa were found to have significantly different niches with respect to both *D* distributions (*P* < 0.05). Plots for all comparisons are available in Appendix 1. Figure prepared with Inkscape 1.2 (Inkscape Project, 2022).

**Table 4.**
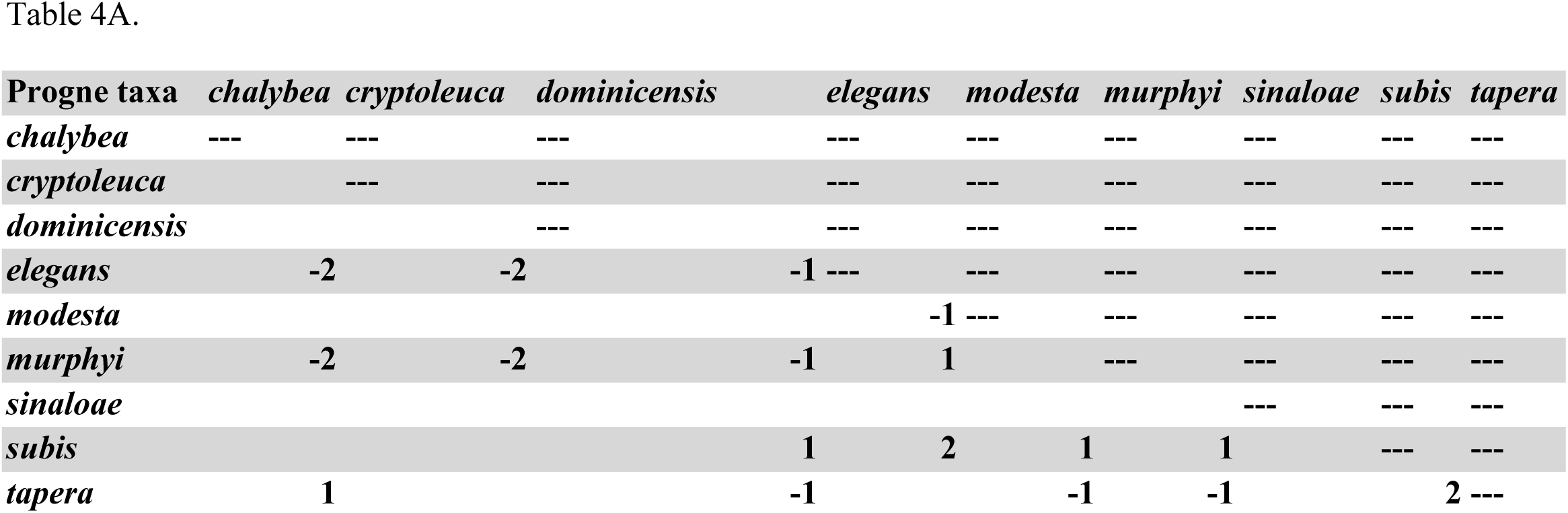
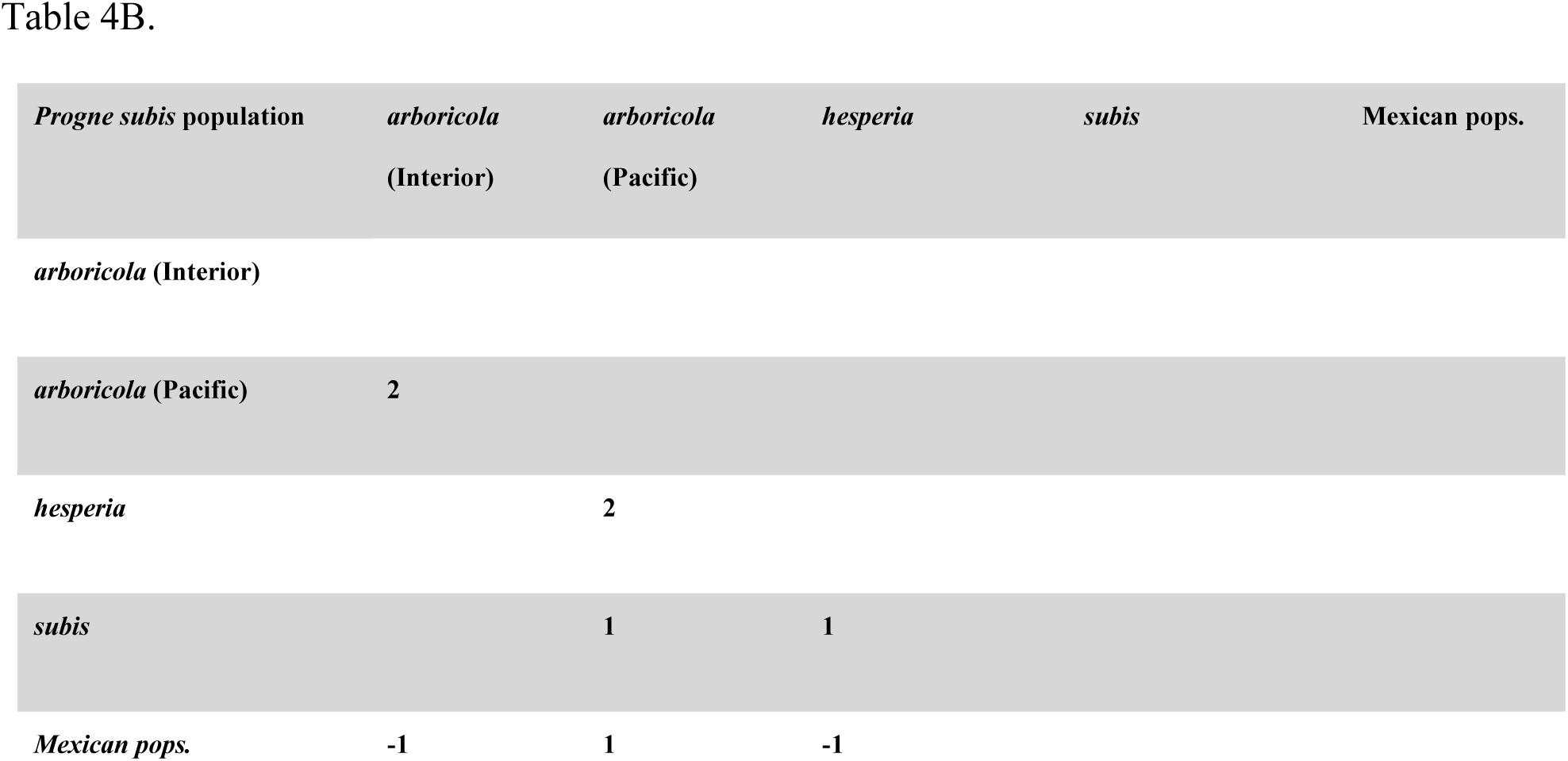
Significance of comparisons of ecological niches using random distributions drawn from each test species. Negative values indicate niche divergence, and positive values extreme conservatism, with only a score of ‘-2’ being considered significant niche divergence. *A*: Interspecific niche comparisons within *Progne*. *B*: Intraspecific niche comparisons within *P. subis*.

### 3.3 Ecological Niche Reconstructions

Reconstructions were created for each variable independently. The ancestral *Progne* was found to have a broad ecological niche in most aspects, though (in some cases) not quite as broad as the most basal extant species, *P. tapera* (Figure 3; see Appendix 1). Reconstructions consistently showed instances of niche contraction in geographically localized taxa, especially among the restricted-range species such as *P. murphyi, P. sinaloae, and P. cryptoleuca*. Niche expansion was observed for several traits, such as Emberger’s Pluviometric Quotient (designed to separate Mediterranean climates), but such expansions were mostly observed in widespread taxa or taxa that breed at high latitudes. Some taxa also experienced niche expansion with respect to their sister species, further indicating flux in the occupied environmental areas within the clade. Wide ranging taxa, especially *P. subis* and *P. chalybea*, demonstrate this niche expansion with respect to their most recent common ancestor with their less widespread sister taxa. Most niches, however, were similar to the broad ancestral niche, or were contained within the space of the broad ancestral niche. Unsurprisingly, some taxa, especially insular taxa, inhabit ecological niches that cannot be characterized completely owing to limited environmental conditions being present within the species’ accessible area.

**Figure 3.**
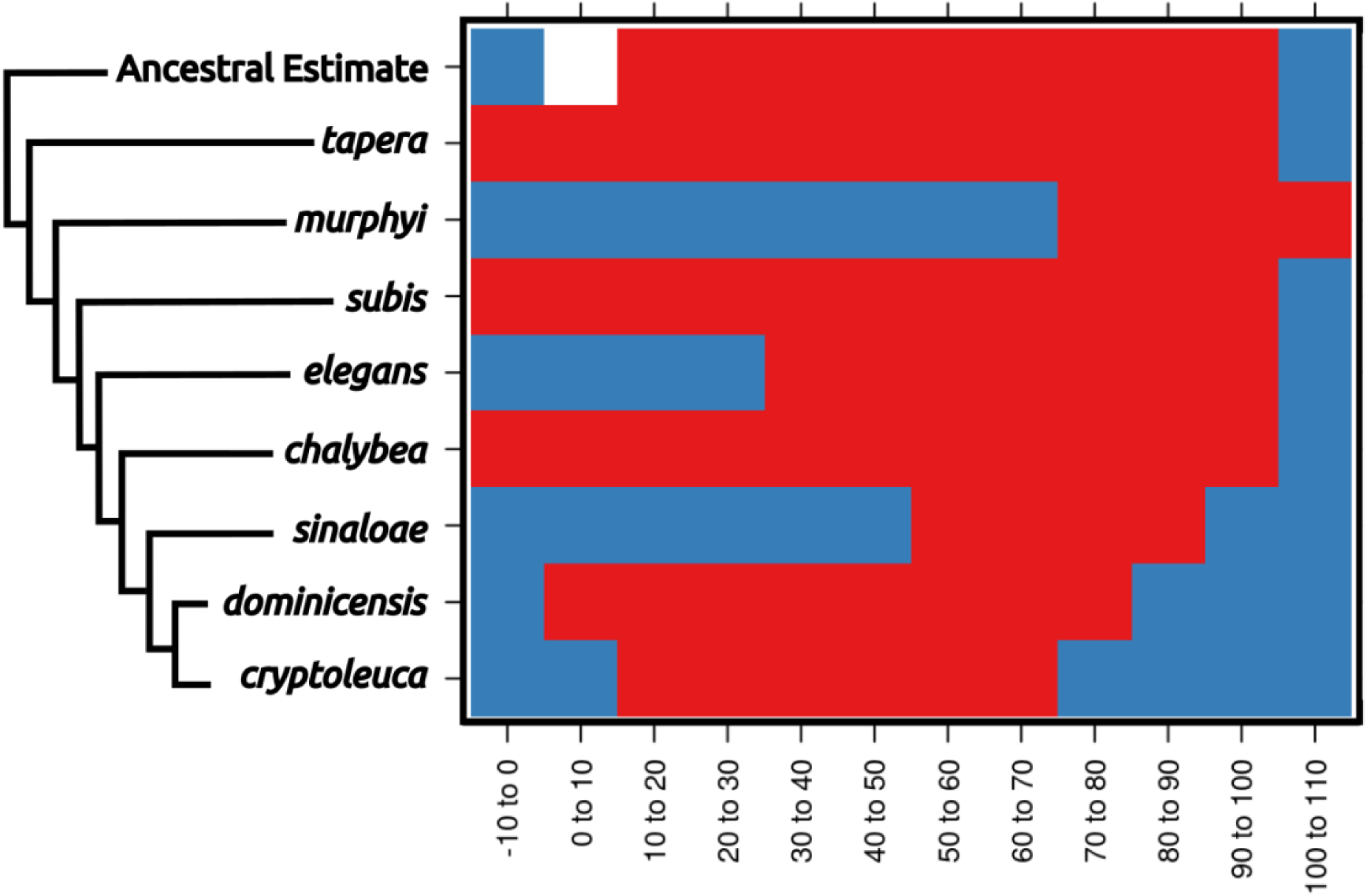
Ecological niche reconstructions for aridity tolerance for the genus *Progne*, with niche ranges shown across the plot as occupied (red), unoccupied (blue), or uncharacterized (white). Note that the historic niche appears to be rather large and encompassed most of the niche now occupied by the entire genus. Widespread modern taxa, such as *P. tapera*, *P. subis*, and *P. chalybea*, have large niches similar to the ancestral state, while restricted range taxa have evolved into narrower niche spaces (with *P. murphyi* even showing evidence for niche expansion), and some of these taxa would appear to have niche divergence if not compared within the larger framework of the genus *Progne*. A scaleless phylogeny based on Brown (2019) is shown for reference, with the basal node shown as the “Ancestral Estimate”. Figure prepared with Inkscape 1.2 (Inkscape Project, 2022) and R package *rasterVis* (Perpiñán & Hijmans, 2023).

## 4 Discussion

I found widespread support for niche conservatism within *Progne*. Historical reconstructions demonstrate that members of *Progne* are largely overlapping in ecological space both with their congeners and the presumed niche of their ancestral taxon. Thus, current niche similarities are not the result of ecological niche convergence, and observed divergences appear to be the result of niche partitioning wherein species niches contract with respect to their ancestral state. The most similar ecological niches were found among basal martins with broad distributions, namely between *P. subis* and both *P. tapera* and *P. elegans*.

I found only a few instances of ecological niche divergence overall, with these divergences being between non-sister taxa. Two of the incidences of niche divergence were between *P. elegans*, the outgroup of the ‘white-breasted’ martin clade, and species found in the same clade (*P. chalybea* and *P. cryptoleuca*). This group includes multiple morphologically conservative species that are not always identifiable in the field, but that possess largely discrete breeding ranges in diverse habitats, ranging from mid-elevation montane forests to coastal scrub, palms, and urban areas (Fang & Schulenberg, 2020; García-Lau & Turner, 2021; Perlut & Williams, 2021).

### 4.1 Ecological niche conservatism in Progne

I was unable to reject the null hypothesis of niche conservatism for most of *Progne*, providing further evidence that organisms are unlikely to have rapid niche shifts while diversifying (Peterson, Soberón & Sanchez-Cordero, 1999). Evolutionary reconstructions indicate that the ancestral *Progne* was a South American species with a broad ecological niche, and that it was at least partially migratory (Brown, 2019), similar to the ecology of the extant *Progne tapera*. The next basal groups included *P. subis*, a long-distance North American migrant, and the range-restricted species pair *P. murphyi* and *P. modesta*. It appears that *P. subis* is descended from a South American species (Brown, 2019), perhaps similar to the recent North-to-South Hemisphere colonization of Barn Swallows *Hirundo rustica* in Argentina (Martínez, 1983). Similar north-to-south shifts and geographic isolation appear to have led to diversification within the ‘white-breasted’ martins as well, resulting in three allopatric species breeding across the North American and Caribbean tropics (Brown, 2019). Such events demonstrate the importance in geographic isolation for driving diversification within *Progne*, and support the idea of *Progne* undergoing a ‘geographic radiation’ of speciating while maintaining similar ecological niches among the descendant species (Peterson, Soberón & Sanchez-Cordero, 1999; Simões et al., 2016).

Ecological niche differences exist among a few extant *Progne*, but these appear to be the result of ecological niche partitioning and not true ecological niche shifts. As species have colonized isolated regions and have continued to adapt regionally, they have experienced niche contractions, possibly as a result of specializing to conditions in these regions, or moderate niche shifts, wherein they have expanded their ecological tolerance within their specific geographic distributions. The former is demonstrated well by *Progne murphyi*, a species restricted to the coast of Peru. The restricted ecological niche of P. murphyi is reminiscent of taxon cycles in the Caribbean: species diversify, and descendant species become more restricted (and sometimes more ecologically specialized) through time (Ricklefs & Cox, 1972; Engler et al., 2021). Results within the genus Progne indicate that tests of ecological niche divergence should try to account for ancestral niche states to understand whether observed niche evolution is novel or reflective of a different evolutionary process, such as partitioning of the ancestral state.

*Progne subis* appears to be a microcosm of these phenomena, with the species as a whole occupying a broad ecological niche, with individual populations evolving for specific environmental conditions within the overall niche space. Within *P. subis*, evidence for niche divergence is lacking, with results reflecting a continuum of variation between descendant populations partitioning environments than any true ecological shift. In southwestern North America, cactus-nesting *P. s. hesperia* are found in close geographic proximity with montane *P. s. arboricola,* yet I found ecological niche overlap between these taxa that differ drastically with respect to habitat preference. These populations have niches within the broader *Progne* subis niche reminiscent of other extant species occupying subsets of the broader *Progne* niche elsewhere (*e.g., P. murphyi* and *P. sinaloae*), further supporting the notion of niche partitioning and niche specialization across a geographic mosaic, as opposed to significant ecological niche evolution leading to large ecological shifts. Diversification via niche partitioning of a broader ancestral state may be more common than is realized in continental taxa, given the propensity of groups like *Zosterops* to undergo taxon cycles in montane regions (Ricklefs & Cox, 1972; Melo, Warren & Jones, 2011; Pearson & Turner, 2017; Engler et al., 2021), and the propensity for related populations to partition and specialize on different food sources across space and time (Benkman et al., 2009; Cenzer, 2016; Alonso et al., 2020).

### 4.2 Assessments of niche in geographically widespread groups

Assessments of ecological niches are still frequently performed across political boundaries or study area boundaries, and thus do not account for the bias introduced by including sites and associated environments not accessible to the study taxa over relevant time periods (Soberón & Peterson, 2005; Barve et al., 2011; Owens et al., 2013; Peterson & Anamza, 2015; Cooper & Soberón, 2018; Song et al., 2020). These oversights can bias niche estimates and therefore bias distribution models in current environments. These issues can also compound when comparing ecological niches through time to understand their evolution (Saupe et al., 2018).

Another less discussed oversight, however, is that of population-level variation within species. What constitutes a species can be contentious (Watson, 2005), so what is recognized as a species in the literature can vary greatly between taxonomic authorities (Barrowclough et al., 2016; Garnett & Christidis, 2017; Raposo et al., 2017). The effects of these oversights are easily missed in studies focusing on ‘species-level’ diversification. I focused on sets of populations of *P. subis*, and results illustrated the processes that are occurring on a more unitary, fine-scale basis within this one species.

## 5 Conclusions and Future Directions

Future research on *Progne* should focus on understanding relationships between populations across ranges of species (particularly *P. subis*) and in clarifying breeding and non-breeding distributions within the genus. Several species and populations are poorly known, most notably populations of *P. subis* and *P. sinaloae* in western Mexico. Ecological analyses such as those developed here are useful for helping identify ecological specialization between closely related taxa, even when those taxa occupy portions of the broad ancestral niche, and can guide future efforts to plan sampling for phylogeographic analyses. Ecological niche analyses can also focus more on the temporality of ecological niches, specifically considering nest sites, wintering sites, and the seasonal occupancy of each species or population, to understand environmental conditions necessary for species survival and to understand how niches are or are not conserved through annual cycles (Nakazawa et al., 2004; Peterson et al., 2005).

Ecological niche divergence is not a driver of or an inevitable consequence of diversification, and geographically widespread radiations can exhibit niche conservatism. *Progne* martins provide an excellent case study of cryptic diversification driven by habitat specialization, behavioral differences, and allopatry. Species pairs that occupy small, specialized ranges or different latitudinal areas can show ecological niche differences from related taxa while still occupying portions of the ancestral ecological niche. Within the ecologically diverse *P. subis*, niche conservatism between described subspecies appears pervasive, notwithstanding divergence in habitat preferences and behavior. These results highlight how niche divergence can be decoupled from diversification, and highlight the need for extensive geographic sampling when studying gene flow and variation within taxa.

## 6 Acknowledgments

I would like to dedicate this paper to Richard G. Levad, whose personal mentorship helped inspire me to study birds and whose work shed light on Colorado’s most enigmatic birds, including *Progne subis* populations there. I would like to thank Kim Potter, Jason Beason, Glenn P. Giroir, and Carolyn Gunn for further instilling in me an interest in *Progne* martins. A. Townsend Peterson provided critical comments on this manuscript. Clare E. Brown provided comments regarding the project and enabled this research with her graduate work. Additional phylogenetic assistance was provided by Subir Shakya and Fred Sheldon. Coding advice and assistance were provided by Marlon Cobos, Fernando Machado-Stredel, Jorge Soberón, and Joseph D. Manthey.

## 7 Data Accessibility

All code are available in Appendix 1 and in the Supplemental Materials; both are available under a GNU General Public License v3.0 and are accessible from my GitHub page at https://github.com/jacobccooper/progne_niche_evolution.

## 8 Supplemental Materials

All code and additional figures can be found in Appendix 1 and in the Supplemental Materials.

## References

Allen RW, Nice MM. 1952. A study of the breeding biology of the Purple Martin (Progne subis). The American Midland Naturalist 47:606–665. DOI: 10.2307/2422034.

Alonso D, Fernández-Eslava B, Edelaar P, Arizaga J. 2020. Morphological divergence among Spanish Common Crossbill populations and adaptations to different pine species. Ibis 162:1279–1291. DOI: 10.1111/ibi.12835.

Baker AJ, Greenslade AD, Darling LM, Finlay JC. 2008. High genetic diversity in the blue-listed British Columbia population of the Purple Martin maintained by multiple sources of immigrants. Conservation Genetics 9:495–505. DOI: 10.1007/s10592-007-9358-3.

Barrowclough GF, Cracraft J, Klicka J, Zink RM. 2016. How many kinds of birds are there and why does it matter? PLoS ONE 11:e0166307. DOI: 10.1371/journal.pone.0166307.

Barve N, Barve V, Jiménez-Valverde A, Lira-Noriega A, Maher SP, Peterson AT, Soberón J, Villalobos F. 2011. The crucial role of the accessible area in ecological niche modeling and species distribution modeling. Ecological Modelling 222:1810–1819. DOI: 10.1016/j.ecolmodel.2011.02.011.

Benkman CW, Smith JW, Keenan PC, Parchman TL, Santisteban L. 2009. A new species of the Red Crossbill (Fringillidae: Loxia) from Idaho. The Condor 111:169–176. DOI: 10.1525/cond.2009.080042.

Billerman SM, Keeney BK, Rodewald PG, Schulenberg TS (eds.). 2020. Birds of the World. Ithaca, New York: Cornell Laboratory of Ornithology.

Bivand R, Keitt T, Rowlingson B. 2022. rgdal: Bindings for the Geospatial Data Abstraction Library. R package version 1.5–32.

Bivand RS, Lewin-Koh N. 2022. maptools: Tools for handling spatial objects, R package version 1.1–4.

Borregaard MK, Rahbek C, Fjeldsaa J, Parra JL, Whittaker RJ, Graham CH. 2014. Node-based analysis of species distributions. Methods in Ecology and Evolution 5(11):1225–1235. DOI: 10.1111/2041-210X.12283.

Brown CE. 2019. Phylogeny and evolution of swallows (Hirundinidae) with a transcriptomic perspective on seasonal migration. Doctoral Dissertation Thesis. Baton Rouge, Louisiana: Louisiana State University.

Brown CR, Airola DA, Tarof S. 2021. Purple Martin (Progne subis), version 2.0. Birds of the World. DOI: 10.2173/bow.purmar.02.

Cenzer ML. 2016. Adaptation to an invasive host is driving the loss of a native ecotype. Evolution 70:2296–2307. DOI: 10.1111/evo.13023.

Cicero C, Pyle P, Patten MA. 2020. Juniper Titmouse (Baeolophus ridgwayi), version 1.0. Birds of the World. DOI: 10.2173/bow.juntit1.01.

Cobos ME, Cheng Y, Song G, Lei F, Peterson AT. 2021. New distributional opportunities with niche innovation in Eurasian snowfinches. Journal of Avian Biology 52. DOI: 10.1111/jav.02868.

Cobos ME, Osorio-Olvera L, Soberon J, Peterson AT, Barve V, Barve N. 2022. ellipsenm: Ecological Niche’s Characterizations Using Ellipsoids.

Cobos ME, Owens HL, Peterson AT. 2020. nichevol: Tools for ecological niche evolution assessment considering uncertainty. R package version 0.1.19.

Comte L, Cucherousset J, Olden JD. 2017. Global test of Eltonian niche conservatism of nonnative freshwater fish species between their native and introduced ranges. Ecography 40:384–392. DOI: 10.1111/ecog.02007.

Cooper JC, Maddox JD, McKague K, Bates JM. 2021. Multiple lines of evidence indicate ongoing allopatric and parapatric diversification in an Afromontane sunbird (Cinnyris reichenowi). Ornithology 138:ukaa081. DOI: 10.1093/ornithology/ukaa081.

Cooper JC, Soberón J. 2018. Creating individual accessible area hypotheses improves stacked species distribution model performance. Global Ecology and Biogeography 27:156–165. DOI: 10.1111/geb.12678.

Corman TE. 2005. Purple Martin. Arizona Breeding Bird Atlas:370–371.

Cuervo PF, Flores FS, Venzal JM, Nava S. 2021. Niche divergence among closely related taxa provides insight on evolutionary patterns of ticks. Journal of Biogeography 48:2865– 2876. DOI: 10.1111/jbi.14245.

Delignette-Muller ML, Dutang C. 2015. fitdistrplus: An R package for fitting distributions. Journal of Statistical Software 64:1–34. DOI: 10.18637/jss.v064.i04.

Dowle M, Srinivasan A. 2019. data.table: Extension of “data.frame”. R package version 1.12.6.

Dubay SG, Witt CC. 2014. Differential high-altitude adaptation and restricted gene flow across a mid-elevation hybrid zone in Andean tit-tyrant flycatchers. Molecular Ecology 23:3551–3565. DOI: 10.1111/mec.12836.

eBird. 2012. eBird: An online database of bird distribution and abundance [web application]. eBird, Ithaca, New York.

van Els P, Herrera-Alsina L, Pigot AL, Etienne RS. 2021. Evolutionary dynamics of the elevational diversity gradient in passerine birds. Nature Ecology & Evolution 5:1259– 1265. DOI: 10.1038/s41559-021-01515-y.

Endler JA. 1977. Geographic variation, speciation, and clines. Princeton, N.J: Princeton University Press.

Engler JO, Lawrie Y, Cabral JS, Lens L. 2021. Niche evolution reveals disparate signatures of speciation in the ‘great speciator’ (white-eyes, Aves: Zosterops). Journal of Biogeography 48:1981–1993. DOI: 10.1111/jbi.14128.

Fang ED, Schulenberg TS. 2020. Sinaloa Martin (Progne sinaloae), version 1.0. Birds of the World. DOI: 10.2173/bow.sinmar1.01.

García-Lau I, Bani Assadi S, Kent G, González A, Rodríguez-Ochoa A, Jiménez A, Acosta M, Mugica L, Meyer K. 2021. Tracking Cuban Martin (Progne cryptoleuca) migration to wintering location and back using geolocators: solving a mystery. Ornithology Research 29:106–112. DOI: 10.1007/s43388-021-00057-y.

García-Lau I, Turner A. 2021. Cuban Martin (Progne cryptoleuca), version 2.0. Birds of the World. DOI: 10.2173/bow.cubmar.02.

García-Navas V, Westerman M. 2018. Niche conservatism and phylogenetic clustering in a tribe of arid-adapted marsupial mice, the Sminthopsini. Journal of Evolutionary Biology 31:1204–1215. DOI: 10.1111/jeb.13297.

Garnett ST, Christidis L. 2017. Taxonomy anarchy hampers conservation. Nature 546:25–27. DOI: 10.1038/546025a.

Global Biodiversity Information Facility. 2022. GBIF Occurrence Download for Progne. DOI: 10.15468/dl.btsx3g.

Glor RE, Warren D. 2011. Testing ecological explanations for biogeographic boundaries. Evolution 65:673–683. DOI: 10.1111/j.1558-5646.2010.01177.x.

Helms JA, Godfrey AP, Ames T, Bridge ES. 2016. Predator foraging altitudes reveal the structure of aerial insect communities. Scientific Reports 6:28670. DOI: 10.1038/srep28670.

Hijmans RJ. 2022. raster: Geographic Data Analysis and Modeling. R package version 3.5–15.

Hijmans RJ, Phillips SJ, Leathwick J, Elith J. 2021. dismo: Species distribution modeling. R package version 1.3–5.

Hu J, Jiang Z, Chen J, Qiao H. 2015. Niche divergence accelerates evolution in Asian endemic Procapra gazelles. Scientific Reports 5:10069. DOI: 10.1038/srep10069.

Hubbard JP. 1969. The relationships and evolution of the Dendroica coronata complex. The Auk 86:393–432.

Inkscape Project. 2022. Inkscape.

Kassambara A, Mundt F. 2020. factoextra: Extract and visualize the results of multivariate data analyses. R package version 1.0.7.

Kennedy JD, Marki PZ, Reeve AH, Blom MPK, Prawiradilaga DM, Haryoko T, Koane B, Kamminga P, Irestedt M, Jønsson KA. 2022. Diversification and community assembly of the world’s largest tropical island. Global Ecology and Biogeography n/a. DOI: 10.1111/geb.13484.

Khaliq I, Fritz SA, Prinzinger R, Pfenninger M, Böhning-Gaese K, Hof C. 2015. Global variation in thermal physiology of birds and mammals: Evidence for phylogenetic niche conservatism only in the tropics. Journal of Biogeography 42:2187–2196. DOI: 10.1111/jbi.12573.

Kozak KH, Wiens JJ. 2006. Does niche conservatism promote speciation? A case study in North American salamanders. Evolution 60:2604. DOI: 10.1554/06-334.1.

Lagasse B. 2020. Gray-breasted Martin (Progne chalybea), version 1.0. Birds of the World. DOI: 10.2173/bow.gybmar.01.

Luo MK. 2020. Peruvian Martin (Progne murphyi), version 1.0. Birds of the World. DOI: 10.2173/bow.permar1.01.

Martínez MM. 1983. Nidificación de Hirundo rustica erythrogaster (Boddaert) en la Argentina (Aves, Hirundinidae). Neotropica 29:83–86.

McCormack JE, Zellmer AJ, Knowles LL. 2009. Does niche divergence accompany allopatric divergence in Aphelocoma jays as predicted under ecological speciation? Insights from tests with niche models. Evolution 65:1–14. DOI: 10.1111/j.1558-5646.2009.00900.x.

Melo M, Warren BH, Jones PJ. 2011. Rapid parallel evolution of aberrant traits in the diversification of the Gulf of Guinea white-eyes (Aves, Zosteropidae). Molecular Ecology 20:4953–4967. DOI: 10.1111/j.1365-294X.2011.05099.x.

Milá B, Smith TB, Wayne RK. 2007. Speciation and rapid phenotypic differentiation in the Yellow-rumped Warbler Dendroica coronata complex. Molecular Ecology 16:159–173. DOI: 10.1111/j.1365-294X.2006.03119.x.

Mostrom AM, Curry RL, Lohr B. 2020. Carolina Chickadee (Poecile carolinensis), version 1.0. Birds of the World. DOI: 10.2173/bow.carchi.01.

Moyle RG, Slikas B, Whittingham LA, Winkler DW, Sheldon FH. 2008. DNA Sequence assessment of phylogenetic relationships among New World martins (Hirundinidae: Progne). The Wilson Journal of Ornithology 120:683–691. DOI: 10.1676/07-131.1.

Nakazawa Y, Peterson AT, Martínez-Meyer E, Navarro-Sigüenza AG. 2004. Seasonal Niches of Nearctic-Neotropical Migratory Birds: Implications for the Evolution of Migration. The Auk 121:610–618. DOI: 10.1093/auk/121.2.610.

Oksanen J, Simpson GL, Blanchet FG, Kindt R, Legendre P, Minchin PR, O’Hara RB, Solymos P, Stevens MHH, Szoecs E, Wagner H, Barbour M, Bedward M, Bolker B, Borcard D, Carvalho G, Chirico M, De Caceres M, Durand S, Evangelista HBA, FitzJohn R, Friendly M, Furneaux B, Hannigan G, Hill MO, Lahti L, McGlinn D, Ouellette M-H, Ribeiro Cunha E, Smith T, Stier A, Ter Braak CJF, Weedon J. 2022. vegan: Community ecology package. R package version 2.6–2.

Owens HL, Campbell LP, Dornak LL, Saupe EE, Barve N, Soberón J, Ingenloff K, Lira-Noriega A, Hensz CM, Myers CE, Peterson AT. 2013. Constraints on interpretation of ecological niche models by limited environmental ranges on calibration areas. Ecological Modelling 263:10–18. DOI: 10.1016/j.ecolmodel.2013.04.011.

Parker TAI, Stotz DF, Fitzpatrick JW. 1996. Ecological and distributional databases. In: Neotropical birds: ecology and conservation. Chicago, Illinois, USA: University of Chicago Press, 113–436.

Pearson DJ, Turner DA. 2017. A taxonomic review of the genus Zosterops in East Africa, with a revised list of species occurring in Kenya, Uganda and Tanzania. Scopus 37:1–13.

Pebesma E. 2018. Simple features for R: Standardized support for spatial vector data. The R Journal 10:439–446.

Pennell MW, Eastman JM, Slater GJ, Brown JW, Uyeda JC, FitzJohn RG, Alfaro ME, Harmon LJ. 2014. geiger v2.0: An expanded suite of methods for fitting macroevolutionary models to phylogenetic trees. Bioinformatics 30:2216–2218.

Perlut NG, Klak TC, Rakhimberdiev E. 2017. Geolocator data reveal the migration route and wintering location of a Caribbean Martin (Progne dominicensis). The Wilson Journal of Ornithology 129:605–610. DOI: 10.1676/16-142.1.

Perlut NG, Williams NR. 2021. Caribbean Martin (Progne dominicensis), version 2.0. Birds of the World. DOI: 10.2173/bow.carmar1.02.

Perpiñán O, Hijmans R. 2023. rasterVis. R package version 0.51.5.

Peterson AT. 2011. Ecological niche conservatism: A time-structured review of evidence. Journal of Biogeography 38:817–827. DOI: DOI 10.1111/j.1365-2699.2010.02456.x.

Peterson AT, Anamza T. 2015. Ecological niches and present and historical geographic distributions of species: A 15-year review of frameworks, results, pitfalls, and promises. Folia Zoologica 64:207–217.

Peterson AT, Martínez-Campos C, Nakazawa Y, Martínez-Meyer E. 2005. Time-specific ecological niche modeling predicts spatial dynamics of vector insects and human dengue cases. Transactions of The Royal Society of Tropical Medicine and Hygiene 99:647–655. DOI: 10.1016/j.trstmh.2005.02.004.

Peterson AT, Soberón J, Sanchez-Cordero V. 1999. Conservatism of ecological niches in evolutionary time. Science 285:1265–1267. DOI: 10.1126/science.285.5431.1265.

Pistorius A. 1975. Ground-nesting Purple Martins. The Auk 92:814–815.

Prigogine A. 1987. Disjunctions of montane forest birds in the Afrotropical Region. Bonner zoologische Beiträge 38:195–207.

Quigley DTG. 2018. An account of Purple Martin Progne subis records from Ireland and Britain during the 19th century. Irish Birds 11:49–56.

R Core Team. 2022. R: A language and environment for statistical computing. Vienna, Austria: R Foundation for Statistical Computing.

Raposo MA, Stopiglia R, Brito GRR, Bockmann FA, Kirwan GM, Gayon J, Dubois A. 2017. What really hampers taxonomy and conservation? A riposte to Garnett and Christidis (2017). Zootaxa 4317:179–184. DOI: 10.11646/zootaxa.4317.1.10.

Ricklefs RE, Cox GW. 1972. Taxon cycles in the West Indian avifauna. The American Naturalist 106:195–219. DOI: 10.1086/282762.

Roper E. 2020. Galapagos Martin (Progne modesta), version 1.0. Birds of the World. DOI: 10.2173/bow.galmar1.01.

Şahin MK, Candan K, Yildirim Caynak E, Kumlutaş Y, Ilgaz Ç. 2021. Ecological niche divergence contributes species differentiation in worm lizards (Blanus sp.) (Squamata: Amphisbaenia: Blanidae) in Mediterranean part of Anatolian Peninsula and the Levantine region. Biologia 76:525–532. DOI: 10.2478/s11756-020-00548-1.

Saupe EE, Barve N, Owens HL, Cooper JC, Hosner PA, Peterson AT. 2018. Reconstructing ecological niche evolution when niches are incompletely characterized. Systematic Biology 67:428–438. DOI: 10.1093/sysbio/syx084.

Seddon N, Tobias JA. 2007. Song divergence at the edge of Amazonia: An empirical test of the peripatric speciation model. Biological Journal of the Linnean Society 90:173–188. DOI: 10.1111/j.1095-8312.2007.00753.x.

Siddiqui R. 2017. A description and exploration of the potential causes of intra-tropical migration in the Neotropical bird “Purple Martin” (Progne subis). Revue YOUR Review (York Online Undergraduate Research) 3:159–159.

Simões M, Breitkreuz L, Alvarado M, Baca S, Cooper JC, Heins L, Herzog K, Lieberman BS. 2016. The evolving theory of evolutionary radiations. Trends in Ecology & Evolution 31:27–34. DOI: 10.1016/j.tree.2015.10.007.

Soberón J, Peterson AT. 2005. Interpretation of models of fundamental ecological niches and species’ distributional areas. Biodiversity Informatics 2:1–10. DOI: 10.17161/bi.v2i0.4.

Song G, Zhang R, Machado-Stredel F, Alström P, Johansson US, Irestedt M, Mays Jr. HL, McKay BD, Nishiumi I, Cheng Y, Qu Y, Ericson PGP, Fjeldså J, Peterson AT, Lei F. 2020. Great journey of Great Tits (Parus major group): Origin, diversification and historical demographics of a broadly distributed bird lineage. Journal of Biogeography 47:1585–1598. DOI: 10.1111/jbi.13863.

South A. 2017. rnaturalearth: World map data from Natural Earth. R package.

South A. 2022. rnaturalearthhires: High resolution world vector map data from Natural Earth used in rnaturalearth. R package.

Title PO, Bemmels JB. 2018. ENVIREM: An expanded set of bioclimatic and topographic variables increases flexibility and improves performance of ecological niche modeling. Ecography 41:291–307. DOI: 10.1111/oik.02629.

Turner A. 2020. Southern Martin (Progne elegans), version 1.0. Birds of the World. DOI: 10.2173/bow.soumar.01.

Vavrek MJ. 2011. Fossil: Palaoecological and palaeogeographical analysis tools. Palaeontologia Electronica 14.

Venables WN, Ripley BD. 2002. Modern Applied Statistics with S. New York: Springer. DOI: 10.1198/tech.2003.s33.

Vrba ES. 1993. Turnover-pulses, the Red Queen, and related topics. American Journal of Science 293 A:418–452. DOI: 10.2475/ajs.293.A.418.

Warren DL, Glor RE, Turelli M. 2008. Environmental niche equivalency versus conservatism: Quantitative approaches to niche evolution. Evolution 62:2868–2883. DOI: 10.1111/j.1558-5646.2008.00482.x.

Watson DM. 2005. Diagnosable versus distinct: Evaluating species limits in birds. BioScience 55:60–68. DOI: 10.1641/0006-3568(2005)055[0060:DVDESL]2.0.CO;2.

Wickham H, Averick M, Bryan J, Chang W, McGowan LD, François R, Grolemund G, Hayes A, Henry L, Hester J, Kuhn M, Pedersen TL, Miller E, Bache SM, Müller K, Ooms J, Robinson D, Seidel DP, Spinu V, Takahashi K, Vaughan D, Wilke C, Woo K, Yutani H. 2019. Welcome to the tidyverse. Journal of Open Source Software 4:1686. DOI: 10.21105/joss.01686.

